# Fast and accurate large multiple sequence alignments using root-to-leave regressive computation

**DOI:** 10.1101/490235

**Authors:** Edgar Garriga, Paolo Di Tommaso, Cedrik Magis, Ionas Erb, Hafid Laayouni, Fyodor Kondrashov, Evan Floden, Cedric Notredame

## Abstract

Inferences derived from large multiple alignments of biological sequences are critical to many areas of biology, including evolution, genomics, biochemistry, and structural biology. However, the complexity of the alignment problem imposes the use of approximate solutions. The most common is the progressive algorithm, which starts by aligning the most similar sequences, incorporating the remaining ones following the order imposed by a guide-tree. We developed and validated on protein sequences a regressive algorithm that works the other way around, aligning first the most dissimilar sequences. Our algorithm produces more accurate alignments than non-regressive methods, especially on datasets larger than 10,000 sequences. By design, it can run any existing alignment method in linear time thus allowing the scale-up required for extremely large genomic analyses.

**One Sentence Summary:** Initiating alignments with the most dissimilar sequences allows slow and accurate methods to be used on large datasets

## Main Text

Structural and evolutionary predictions are improved when using accurate large Multiple Sequence Alignments (MSAs) featuring thousands of sequences (*1*, *2*). Until the first benchmarking of large-scale MSAs, analyses made on smaller datasets had suggested that scale-up would automatically lead to increased accuracy (*3*). However, all things being equal, alignments with over a thousand sequences are less accurate than their smaller counterparts (*4*). Under the most common multiple alignment procedure (*5*), small MSAs are built in parallel and progressively merged into increasingly large intermediate MSAs, following the order of a guide-tree. This sometimes results in extremely gapped MSAs (*6*) that have been speculated to account for the accuracy drop on large datasets (*7*). Recent attempts to address this problem have included SATé (*8*) and its follow-up PASTA (*9*), a progressive algorithm in which the guide-tree is split into subsets that are independently aligned and later merged. This divide and conquer strategy allows computationally intensive methods to be deployed on large datasets but does not alleviate the challenge of merging very large intermediate MSAs. More recent alternatives include the MSA algorithms UPP (*10*) and MAFFT-Sparsecore (Sparsecore) (*11*). Both rely on selecting a subset of sequences - the seeds - and turning them into a Hidden Markov Model (HMM) using either PASTA or the slow/accurate version of MAFFT. The HMM is then used to incorporate all the remaining sequences one by one. The downside of this approach is that the seed sequences can be insufficiently diverse and therefore preclude the accurate alignment of distantly related homologous to the seed HMM. We propose a regressive algorithm that addresses this problem by combining the benefits of a progressive approach when incorporating distant homologues with the improved accuracy of seeded methods. This synthesis was achieved by fulfilling two simple constraints: the splitting of the sequences across sub-MSAs each containing a limited number of sequences and their combination into a final MSA without the requirement of an alignment procedure. Thus, the main difference between our approach and existing ones lies in the order in which sequences are aligned, starting with the most diverse.

Given M sequences the sub-MSAs are collected as follows. A clustering algorithm is used to define N non-overlapping sequence groups - the children. The first sub-MSA - the parent - is computed by selecting a representative sequence from each child group. The same method is then applied onto every child group in which N new representatives are collected and multiply aligned to yield one child sub-MSA for each sequence in the parent. The procedure runs recursively by treating each child as a parent for the next generation until every sequence has been incorporated. The final MSA is produced by merging all the sub-MSAs. The merging of a child with its parent is done without additional alignment thanks to the representative sequence. This sequence, present in both the child and its parent, enables the stacking of the corresponding positions (Fig. 1A). When doing so, insertions occurring within the representative, either in the child or in its parent, are projected as deletions (i.e. gaps) in the other.

**Fig. 1.**
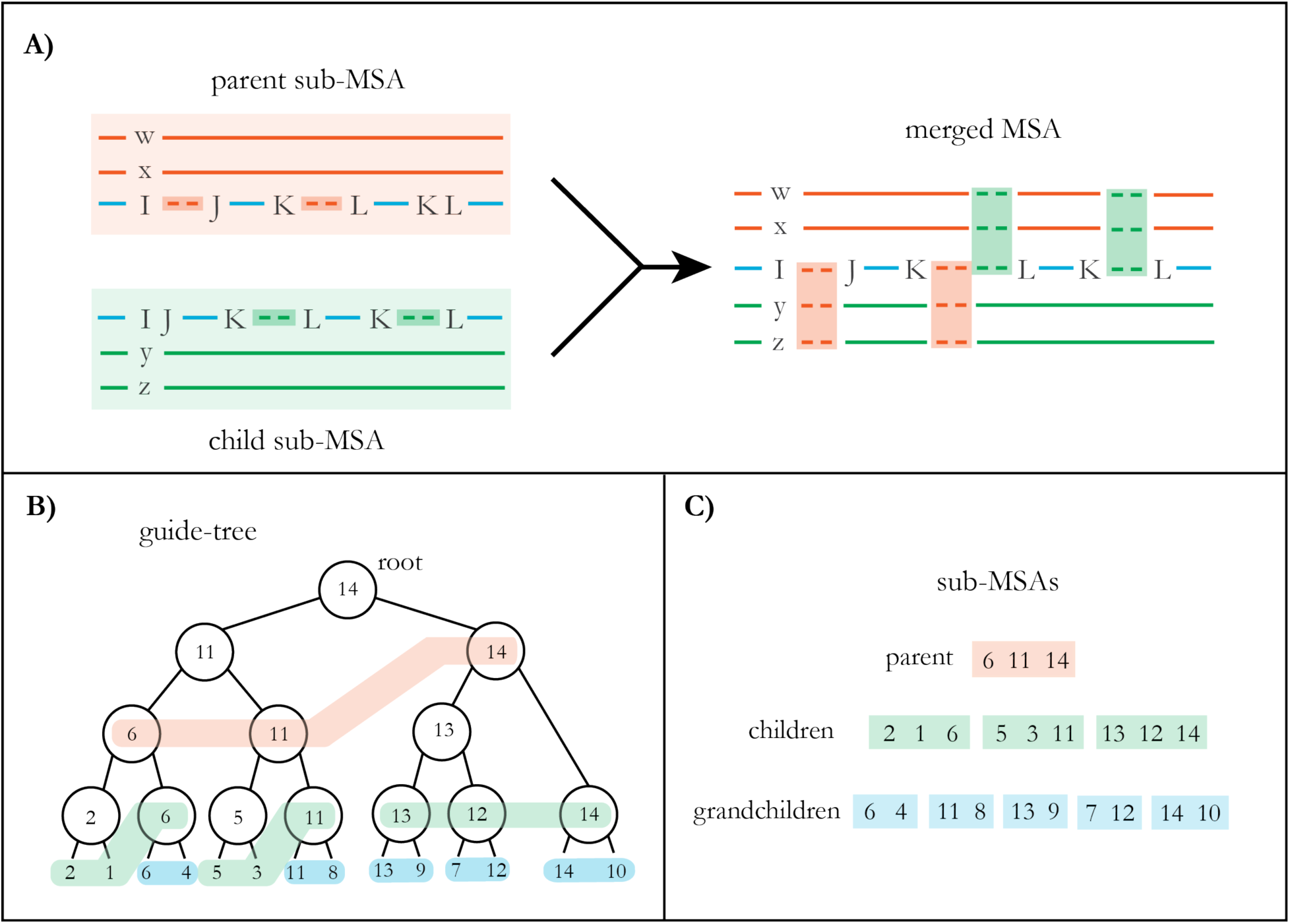
Regressive algorithm overview. (**A**) Parent and children sub-MSAs are merged via their common sequence (blue) whose indels are projected from child to parent (green) and parent to child (red). (**B**) The sub-MSAs are produced after collecting sequences from a binary guide tree with each node labeled with the name of its longest descendant sequence. Sequences are collected by traversing the tree in a breadth-first fashion. Pale red color blocks indicate how the N parent sequences (N=3) are collected by recursively expanding nodes. The same process is then applied to gather the children (green) and the grandchildren (blue). (**C**) In the nine resulting sub-MSAs that are displayed, one should note the presence of a common representative sequence between each child and its parent.

A key step of this recursion is the clustering method and the subsequent selection of the N representative sequences. N was set to 1,000, a figure reported to be the largest number of sequences that can be directly aligned without accuracy loss (*4*). The clusters were estimated from binary guide-trees produced by existing large-scale MSA algorithms such as Clustal Omega (ClustalO) and MAFFT. The use of a binary tree to extract the most diverse sequences was inspired by an existing taxa sampling procedure (*12*). In our implementation, every node gets labeled with the longest sequence among its descendants. Given a fully labeled tree, the sequences of the first parent sub-MSA are collected by the breadth-first traversal of the tree, starting from the root through as many generations as required to collect N sequences (Fig. 1B). Because of the way they are collected along the tree these N first sequences are as diverse as possible. Within the resulting sub-MSA every sequence is either a leaf or the representative of an internal node ready to processed (Fig. 1C).

Our algorithm does not depend on specific alignment or guide-tree methods and therefore lends itself to be combined with any third-party software. This property enabled us to run various alignment software both directly and in combination with the regressive algorithm. A combination involves estimating a guide-tree with an existing method, collecting sequences with the regressive algorithm and then computing the sub-MSAs with an existing MSA algorithm. By doing so we were able to precisely quantify the impact of our algorithm on both accuracy and computational requirements. We used as a benchmark the HomFam protein datasets (*4*) in which sequences with known structures - the references - are embedded among large numbers of homologues. Accuracy is estimated by aligning the large dataset and then comparing the induced alignment of the references with a structure-based alignment of these same references (*13*). We started by benchmarking the ClustalO and MAFFT-FFT-NS-1 (Fftns1) MSA algorithms using two guide-tree methods: ClustalO embedded k-means trees (mBed) (*14*) and MAFFT-PartTree (PartTree) (*15*). These widely adopted software were selected because they support large-scale datasets, are strictly progressive and allow the input and output of binary guide-trees.

In three out of four combinations of guide-tree and MSA algorithms, the regressive combination outperformed the progressive one. When considering the most discriminative measure (total column score, TC, Table 1) on the datasets with over 10,000 sequences, the regressive combination delivered MSAs that were on average 5.13 percentage points more accurate than when computed progressively (39.31 and 34.24 respectively). These differences remained comparable, albeit reduced, when considering the contribution of smaller datasets (Table S1, S2). Within this first set of analysis, the regressive combination of ClustalO with PartTree was the most accurate and on the large datasets it outperformed its progressive counterpart by 15.27 percentage points (42.21 and 26.94 respectively, Wilcoxon p-value <0.001).

**Table 1.**
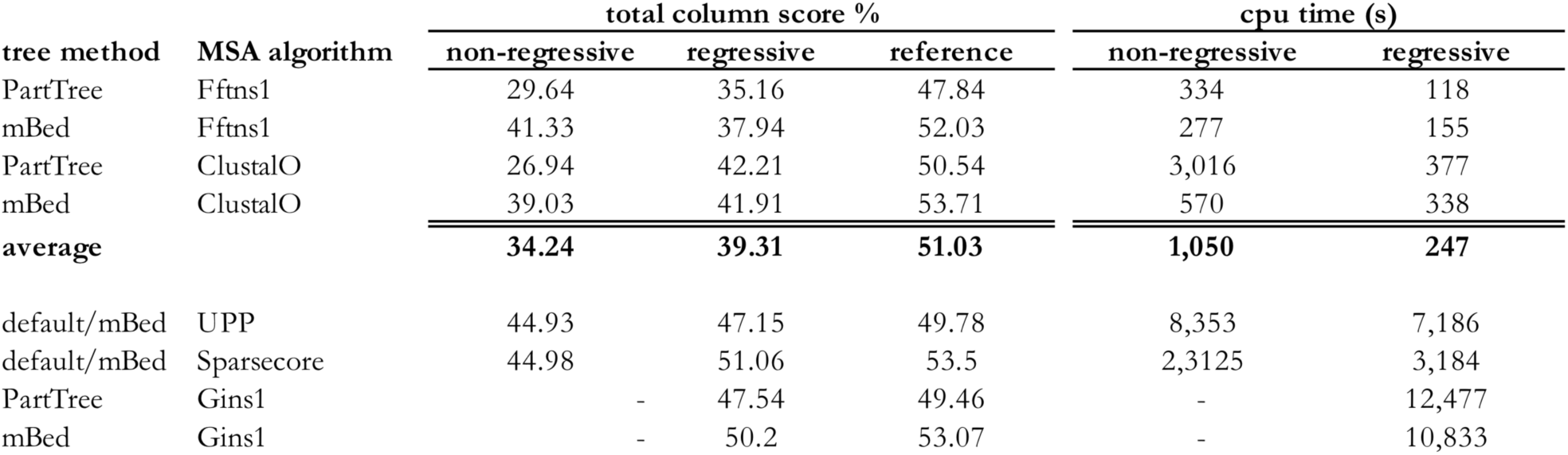
Total Column Score and average CPU time (s) on the 20 HomFam datasets containing over 10,000 sequences.

We also tested the seed-based non-progressive MSA algorithms Sparsecore (*10*) and UPP (*11*). Both were improved when combined with the regressive algorithm. For instance, the regressive combination of Sparsecore with mBed guide-trees yielded the best readouts of this study on the very large alignments, and a clear improvement over the default Sparsecore (TC score 51.07 vs. 44.98, Wilcoxon p-value<0.1). Comparable results were observed when extending this analysis to the Sum-of-Pair (SoP) metrics or to smaller datasets (Table S1, S2). The regressive algorithm is especially suitable for the scale-up of computationally expensive methods. For instance, the consistency-based variant of MAFFT named MAFFT-G-INS-1 (Gins1) (*16*), was among the most accurate small-scale MSA algorithms on the reference sequences. Gins1 cannot, however, be deployed on the HomFam datasets because its computational requirements are cubic with the number of sequences thus restricting it to a few hundred sequences. By combining Gins1 with the regressive algorithm we overcame this limitation and produced the most accurate readouts on datasets larger than 1,000 sequences (Table S1, S2).

We complemented these measures of absolute accuracy with an estimate of accuracy degradation when scaling up. The effect of extra homologous sequences degrading the alignment accuracy of an MSA can be quantified by comparing the small MSAs of the reference sequences alone with their corresponding large-scale datasets. With the default progressive MSA algorithms ClustalO and Fftns1, the large datasets were on average 16.79 percentage points less accurate than when aligning the reference sequences (Table 1, 34.24 and 51.03 respectively) with the trend being amplified on the larger alignments (Fig. 2A). Yet, on this same comparison the regressive combinations were only affected by 11.72 points (Fig. S1). The improved stability of the regressive combination was especially clear when considering Gins1 (Fig. 2A, Fig. S1A) that was merely degraded by 2.87 percentage points thus achieving on the large datasets a level of accuracy close to the one measured on the reference sequences alone (Table 1, 50.20 and 53.07 respectively).

**Fig. 2.**
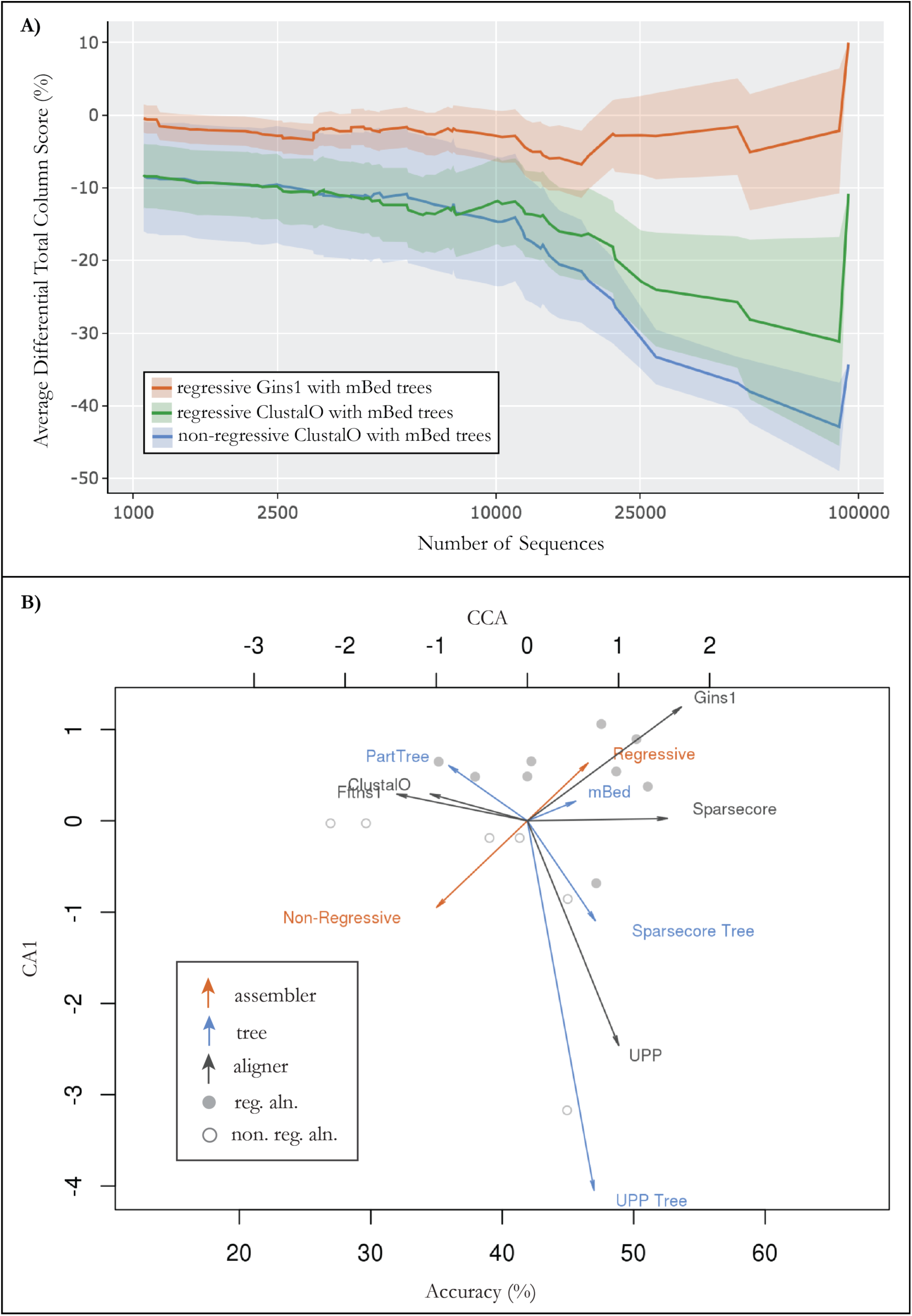
Relative performances of alternative MSA algorithm combinations. (**A**) Average differential accuracy of datasets larger than *Number of Sequences* (horizontal axis). The differences of accuracy are measured between the reference sequence MSAs and their embedded projection in the large datasets. The envelope is the standard deviation. (**B**) In this constrained correspondence analysis (CCA) the first component (horizontal axis, 14.1% of the variance) is constrained to be the total column score accuracy as measured on datasets larger than 10,000. The best unconstrained component (vertical axis) explains 20.8% of the remaining variance. Methods (grey dots with their accuracy on the lower horizontal axis) are categorized by their guide-tree (blue), MSA algorithms (grey) and regressive/non-regressive procedure (red). Vectors indicate the contributions to variance of each category from the three variables. Their projection onto the upper horizontal axis quantifies the contribution to variance of overall accuracy.

Identifying the factors driving accuracy improvement can be challenging considering that each alignment procedure relies on different combinations of algorithmic components (i.e. regressive/non-regressive, tree method, MSA algorithm). For this purpose, we used Constrained Correspondence Analysis (CCA) (*17*), a dimensionality reduction method adapted for categorical variables. When applied to Table 1 data, CCA allowed us to estimate the relative impact of each method’s algorithmic component with respect to a constrained variable - accuracy in this case. As one would expect, the MSA algorithm is the most influential variable with respect to accuracy but CCA confirmed the general benefits of switching from a non-regressive to a regressive combination (Fig. 2B).

When using the same guide-tree for the regressive and non-regressive alignment combinations, improved accuracy comes along with substantially improved computational performance. On average the regressive combinations require about 4-fold less CPU time that their non-regressive equivalent on datasets larger than 10,000 sequences (Table 1). Seeded methods like UPP or Sparsecore appear to benefit less from the regressive deployment with marginal differences in CPU requirements (Fig. 3A). When considering MSA algorithms like ClustalO or Fftns1 that scale linearly with the number of sequences, the improvement yielded by the regressive combination was roughly proportional to the original non-regressive CPU requirements. For instance, in the case of ClustalO using mBed trees, the regressive combination was about twice as fast as the progressive alignment and appeared to have a linear complexity (Fig. 3B). The situation was even more favorable when considering CPU intensive MSA algorithms like Gins1 for which the non-regressive computation had been impossible.

**Fig. 3.**
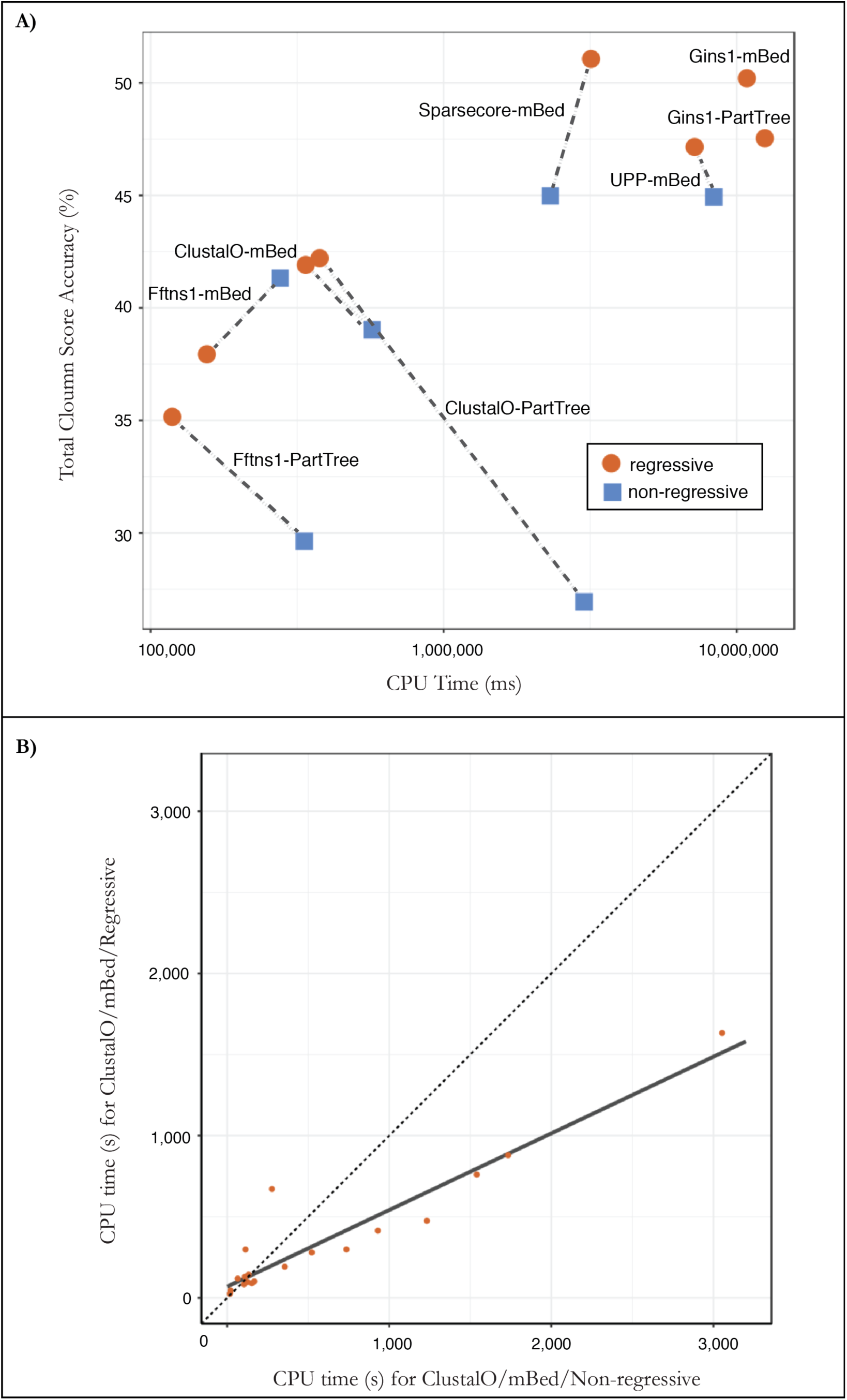
CPU requirements of the regressive algorithm on HomFam datasets containing more than 10,000 sequences. (**A**) The total CPU requirements (horizontal axis) and average total column score accuracies (vertical axis). The corresponding non-regressive (blue square) and regressive (red circles) combinations are connected by a dashed line with the exception of Gins1 for which the non-regressive computation costs are prohibitive. (**B**) Comparison of CPU time requirements for ClustalO using mBed trees using a regressive and a non-regressive procedure on HomFam datasets containing more than 10,000 sequences. A linear regression (grey) was fitted on the resulting graph (R^2^ = 0.89, p-value < 6.9*10^-10^).

The ability to use slow and accurate MSA algorithms in linear time regardless of their original computational complexity is the most important feature of the regressive algorithm. It allows the application of any of these methods - natively - onto extremely large sequence datasets. This linearization is an inherent property of the regressive procedure in which all the sequences are split across sub-MSAs of a bounded size (1,000 sequences). This bounding in size results in a bounded computational cost. Since the total number of sub-MSAs is proportional to the initial number of sequences, the resulting complexity for the full MSA computation is linear. Furthermore, owing to the computational independence of the sub-MSAs, the regressive algorithm turns MSA computation into an embarrassingly parallel problem (*18*).

Our regressive algorithm provides a practical and generic solution to the critical problem of MSA scalability. It is a versatile algorithm that lends itself to further improvements, for instance by exploring the impact of more sophisticated clustering structures, such as k-guide-trees and b-guide-trees or by testing different ways of selecting the representative sequences. The regressive algorithm is nonetheless a mature development framework that will enable a clean break between the improvement of highly accurate small-scale MSA algorithms - like Gins1 - and the design of more efficient large-scale clustering algorithms, like PartTree and mBed. This divide will help potentiate the large body of work carried out in the clustering and alignment communities over the last decades and hopefully speed up the development of new improved methods. Achieving this goal is not optional. There is a Red Queen’s race going on in genomics. It started the day omics’ data growth overtook computing power and it shows no signs of slowing down (*19*).

## Supporting information

## Acknowledgments

We thank Guy Riddihough for revisions and comments on the manuscript and Olivier Gascuel for suggestions.

## Author contributions

C.N. designed and implemented the algorithm, E.F., E.G. and P.D.T carried out the validation, I.E. performed the CCA analysis and all the authors designed the validation procedure and wrote the manuscript.

## Data and materials availability

The regressive alignment algorithm has been implemented in T-Coffee and is available on GitHub for immediate use (https://github.com/cbcrg/tcoffee). All data, analyses and results are available from Zenodo (https://doi.org/10.5281/zenodo.2025847). A GitHub repository containing the Nextflow workflow and Jupyter notebooks to replicate the analysis in full is available at https://github.com/cbcrg/dpa-analysis (release v1.0). See supplementary materials for a written description of methods.

