## Supplementary material for "Fast and accurate large multiple sequence alignments using root-to-leave regressive computation"

**This PDF file includes:**

Materials and Methods
Supplementary Text
Fig. S1
Tables S1 to S2

### Materials and Methods

**Datasets**. The HomFam dataset was downloaded from the Clustal Omega site (*http://www.clustal.org/omega/homfam-20110613-25.tar.gz*) being the last update from the dataset on the 13th June of 2011. This dataset is a group of 94 families with their homologous Pfam sequences from Pfam version 25.

To avoid the fluctuation of the process of embedding the seed sequences into the sequence fasta file, for each family, the exact same input files containing the sequence and reference sequences were used across all analysis and are available in the Git repository under the folder "data/combined_seqs".

**Reproducibility**. A git repository containing all data, analyses performed, and alignments had been archived in the Zenodo storage service and accessible with the following: https://doi.org/10.5281/zenodo.2025847. For convenience, the git repository is also available on GitHub from https://github.com/cbcrg/dpa-analysis with the release tag v1.0. A series of Jupyter notebooks within the notebook directory within this repository contains the necessary commands to replicate the analysis from sequence data through to figures. The pipeline used for generating the guide trees and alignments and performing alignment evaluations was implemented in Nextflow and is available in the same git repository. A Docker image has been created which contains all the pipeline dependencies, thus making it unnecessary to install them in the target environment using either Docker or Singularity. In this distribution, the Docker image used for this computation is available on DockerHub referenced using its uniquely generated sha256 hash:

```
cbcrg/regressive-msa:@sha256:151d41b4f5c359fa20f964156c3cd6709250f3d96ac3da7d9ffc594075e8bede
```

The Dockerfile is also provided in the Git repository to allow for reuse and the addition of new tools.

**Platforms**. Full corpus of results was established on a computational cluster running Scientific Linux release 7.2 with all guide trees, alignments and evaluations carried out within a container using Debian Jessie as the operating system.

**Workflow deployment**. In order to carry out a reproducible deployment of guide tree and alignments, the Nextflow workflow manager was used to deploy containerized versions of the software using Docker or Singularity. Procedures described in the following section can be reproduced by downloading the Nextflow software (*www.nextflow.io*) and running the following command on the Homfam dataset:

```
nextflow run cbcrg/dpa-analysis --seqs="./data/seqs/*.fa" --refs="./data/refs/*.ref' -with-
singularity
```

This command will automatically download the container in which all the aligners and required software have been packaged. The full pipeline will (i) generate large-scale guide trees (PartTree, mBed), (ii) produce, a non-regressive (standard) alignment using either the trees generated before, or the method internal tree when external tree are not supported (iii) use the large-scale guide trees and the aligners to estimate a regressive MSA.

The aligners used are the ones displayed on Table 1: ClustalO, Fftns1, UPP, Gins1 and Sparsecore. The individual tree and MSA generation commands executed by Nextflow can be found in the templates directory of the Git repository and are listed in the Methods section below.

**Software versions**. You can find all the software used in the workflow in the Docker image mention in the section Reproducibility. We also include a list of each tool and their version shown below

*Clustal Omega:* version 1.2.4
http://www.clustal.org/omega/clustal-omega-1.2.4.tar.gz

*MAFFT:* version 7.397 with extensions
http://mafft.cbrc.jp/alignment/software/mafft-7.397-with-extensions-src.tgz

*UPP:* version 4.3.4
http://github.com/smirarab/sepp.git

*T-Coffee*: version dev_@20180723_17:20
checkout: 16617b9d9c393de348e0afae3024eb6c69a4527f
https://github.com/cbcrg/tcoffee.git

**Large scale tree generation.**
mBed and PartTree trees were generated as follows:

*mBed*:

```
clustalo -i ${seqs} --guidetree-out ${id}.${tree_method}.dnd --force
```

*PartTree*:

```
t_coffee -other_pg seq_reformat -in ${seqs} -action +seq2dnd parttree -output newick >>
     ${id}.${tree_method}.dnd
```

(This procedure is a wrapper for the MAFFT-PartTree command that produces a coded binary tree, that must be recoded into a standard newick tree)

**Non-regressive MSA generation**.
The following commands were used to generate non-regressive MSAs.

*Clustal Omega:*

```
clustalo --infile=${seqs}  --guidetree-in=${guide_tree}  --outfmt=fa
     —o ${id}.std.${align_method}.with.${tree_method}.tree.aln
```

*MAFFT-FFT-NS-1:*

```
t_coffee -other_pg seq_reformat -in ${guide_tree} -input newick -in2 ${seqs} -input2 fasta_seq -
action +newick2mafftnewick >> ${id}.mafftnewick

newick2mafft.rb 1.0 ${id}.mafftnewick > ${id}.mafftbinary
```

```
/mafft/bin/mafft --retree 1 --anysymbol --treein ${id}.mafftbinary ${seqs} >
     ${id}.std.${align_method}.with.${tree_method}.tree.aln
```

*MAFFT-G-INS-1:*

```
t_coffee -other_pg seq_reformat -in ${guide_tree} -input newick -in2 ${seqs} -input2 fasta_seq -
action
     +newick2mafftnewick >> ${id}.mafftnewick

newick2mafft.rb 1.0 ${id}.mafftnewick > ${id}.mafftbinary

ginsi --treein ${id}.mafftbinary ${seqs} >${id}.std.${align_method}.with.${tree_method}.tree.aln
```

*MAFFT-Sparsecore:*

```
replace_U.pl ${seqs} // This is a Mafft pre-processing routine

/mafft/bin/mafft-sparsecore.rb -i ${seqs} > ${id}.default.${align_method}.aln
```

*UPP:*

```
replace_U.pl ${seqs}

run_upp.py -s ${seqs}  -m amino -x 1 -o ${id}.default.${align_method}

mv ${id}.default.${align_method}_alignment.fasta ${id}.default.${align_method}.aln
```

**Regressive MSA generation**.
In the following command lines, the flag -dpa_method refers to the alignment method deployed by the T-Coffee. The exact command lines corresponding to these modes are available from the file *lib/util_lib/util_constraints_list.c* in the T-Coffee distribution and are identical to the ones used for the non-regressive method in the previous section.

*Regressive Clustal Omega:*

```
t_coffee -dpa -dpa_method clustalo_msa   -dpa_tree ${guide_tree} -seq ${seqs}  -dpa_nseq
${bucket_size} –outfile ${id}.dpa_${bucket_size}.${align_method}.with.${tree_method}.tree.aln
```

*Regressive MAFFT-FFT-NS-1:*

```
t_coffee -dpa -dpa_method mafftfftns1_msa -dpa_tree ${guide_tree} -seq ${seqs} -dpa_nseq
${bucket_size} –outfile ${id}.dpa_${bucket_size}.${align_method}.with.${tree_method}.tree.aln
```

*Regressive MAFFT-G-INS-1:*

```
t_coffee -dpa -dpa_method mafftginsi_msa -dpa_tree ${guide_tree} -seq ${seqs} -dpa_nseq
${bucket_size} –outfile ${id}.dpa_${bucket_size}.${align_method}.with.${tree_method}.tree.aln
```

*Regressive MAFFT-Sparsecore:*

```
t_coffee -dpa -dpa_method mafftsparsecore_msa -dpa_tree ${guide_tree} -seq ${seqs} -dpa_nseq
${bucket_size} -outfile ${id}.dpa_${bucket_size}.${align_method}.with.${tree_method}.tree.aln
```

*Regressive UPP:*

```
t_coffee -dpa -dpa_method upp_msa -dpa_tree ${guide_tree} -seq ${seqs} -dpa_nseq ${bucket_size} -
outfile ${id}.dpa_${bucket_size}.${align_method}.with.${tree_method}.tree.aln
```

**Benchmarking**. Benchmarking calculations were carried out using the aln_compare component of the T-Coffee package that supports the sums of pairs and the total column score metrics. The metrics can be described as follows:

Sums of pairs (SoP): given every pair of aligned residues in the reference MSA, defined by their index and ignoring gaps, the SoP score is the fraction of these pairs found in the target MSA. When the seed sequences are embedded among homologues, the extra homologues are ignored.

Total Column Score (TC): fraction of columns exactly identical between the target and the reference alignment, while taking into account gap symbols - ignoring the index of their neighbor residues, and residue indexes. This is the most stringent measure currently available to compare alternative MSAs

The commands used for the MSA evaluation using the metrics implemented with the T-Coffee package were executed as follows:

*Sum Of Pairs (SoP):*

```
t_coffee -other_pg aln_compare -al1 ${ref_alignment} -al2 ${test_alignment}.compare_mode sp |
grep -v "seq1" |grep -v '*' | awk '{ print \$4}' ORS="\t" >
"${id}.${align_type}.${bucket_size}.${align_method}.${tree_method}.sp"
```

*Total Column Score (TC):*

```
t_coffee -other_pg aln_compare -al1 ${ref_alignment} -al2 ${test_alignment} -compare_mode tc |
grep -v "seq1" |grep -v '*' | awk '{ print \$4}' ORS="\t" >
"${id}.${align_type}.${bucket_size}.${align_method}.${tree_method}.tc"
```

**Supplementary Methods**

Regressive algorithm pseudocode

The exact implementation is available on the GitHub repository as part of the T-Coffee latest beta distribution (https://github.com/cbcrg/tcoffee). The following pseudo-code provides a simplified overview of a serial version of the algorithm assuming a binary tree in which every node is labeled with the name of the longest sequence among its descendant leaves. The symbol node refers to three-ways nodes in this binary tree where each node has a parent, a left child, a right child and a label, with the exception of leafs that have no child and the root that has no parent. The regressive algorithm traverses the binary tree in a recursive fashion, when doing so, it collects N sequences, multiply align them and merges the resulting sub-MSAs. The code is initiated with the node=root of the labeled tree.

```
node2subMSA (node)
    // expands node into a linked list
    // start->node1->...->nodeN->end
    // Turns the linked list into a parent_MSA
    // recursively expand each parent_MSA sequence into a child_MSA
    // merges every child_MSA within the parent_MSA
    // returns the parent_MSA
    // gets initialized with node2subMSA (root)

    if (node is leaf) return node->seq

    start = declare_node()    //empty start node
    end  = declare node()     //empty end node

    //define the linked list: start->node->end
    node->next = end; end-> prev = node
    node->prev = start; start-> next = node

    n=1           //sequences counter
    nleaf=0       //leaf counter

    //Expand all the nodes, one generation at a time

    while (n<N and !(node==end and n==nleaves))

      //rewind if all the nodes if
      //the current generation has been expanded
      if (node==end)node=start->next

      //replace current node with its left and right children
      //Turn pnode->node->nnode into pnode->left->right->nnode
      if (node is not leaf)
            pnode=node->previous
            nnode=node->next

            pnode->next=node->left
            node->left->prev=pnode
            node->left->next=node->right
            node->right->prev=node->left
            node->left->next=nnode
            nnode->prev=node->left
```

```
            //the sequence list is now one sequence longer
            n=n+1

            //keep advancing one node at a time
            node=nnode

      else
            //keep track of the leaf nodes
            nleaves=nleaves+1

    //Collect all the sequences from start->node1->node2…->end
    node=start->next
    while (node !=end)
      add node->label to SeqList
      node=node->next

    //Send collected sequences to a third party aligner
    parent_MSA=MSA_Algorithm(SeqList)

    //Treat each internal node in parent_MSA as a start node
    //collect each child_MSA and merge it with parent_MSA
    node=start->next
    while (node!=end)
      child_MSA=node2subMSA (node)
      parent_MSA=merge_subMSAs (node->seq, parent_MSA, child_MSA)
      node=node->next

    //parent_MSA contains ALL descendant sequences of node
    //if node is the root, parent_MSA is the final MSA
    return parent_MSA

MSA_Algorithm (SeqList)
    subMSA=system (Run third party MSA Algorithm on SeqList)
    return subMSA

merge_subMSAs (seq, parent_MSA,child_MSA)
    //seq is the representative sequence
    //seq is present in parent_MSA and in child_MSA

    foreach residues i of seq
      parent_gap=number of '-' between i and i+1 in parent_MSA
      child_gap=number of '-' between i and i+1 in child_MSA

      columnP=parent_MSA column containing residue i of seq
      for each sequence in parent_MSA
            Insert child_gap '-' after columnP

      columnC=child_MSA column containing residue i of seq
      for each sequence in child_MSA
            Insert parent_gap '-' after columnC

    //parent_MSA and Child_MSA now have the same length

    Replace seq in parent_MSA with all sequences of child_MSA
    Return parent_MSA
```

**Constrained Correspondence Analysis.** Each MSA is represented in the form of a string of zeros and ones encoding the categorical variables used by the alignment algorithm. These strings consist of three substrings representing the variables (tree method, aligner, assembly method) and have a length corresponding to the number of levels in each variable (number of possible tree methods etc.). Thus, each substring only contains one 1 entry and the rest 0 entries, the entire sequence sums to 3. MSAs thus represented are the rows of an indicator matrix containing zeros and ones that can be analyzed with dimensionality reduction techniques such as multiple correspondence analysis (*17*). In Constrained (a.k.a. Canonical) Correspondence Analysis (CCA), dimensionality reduction is guided by additional information about each observation. In our case, this information is the average accuracy (column score) of the MSA across the 20 largest data sets. It enters the calculation in form of a vector with components corresponding to the MSAs. Roughly, the suitably transformed indicator matrix is projected onto the linear space defined by the accuracy vector and a singular value decomposition is performed. For mathematical details see (17). Calculations were performed using the R package Vegan (*https://cran.r-project.org/package=vegan*). Percent variance explained is obtained dividing the eigenvalue of the respective axis with the sum of all the eigenvalues, multiplied by 100.

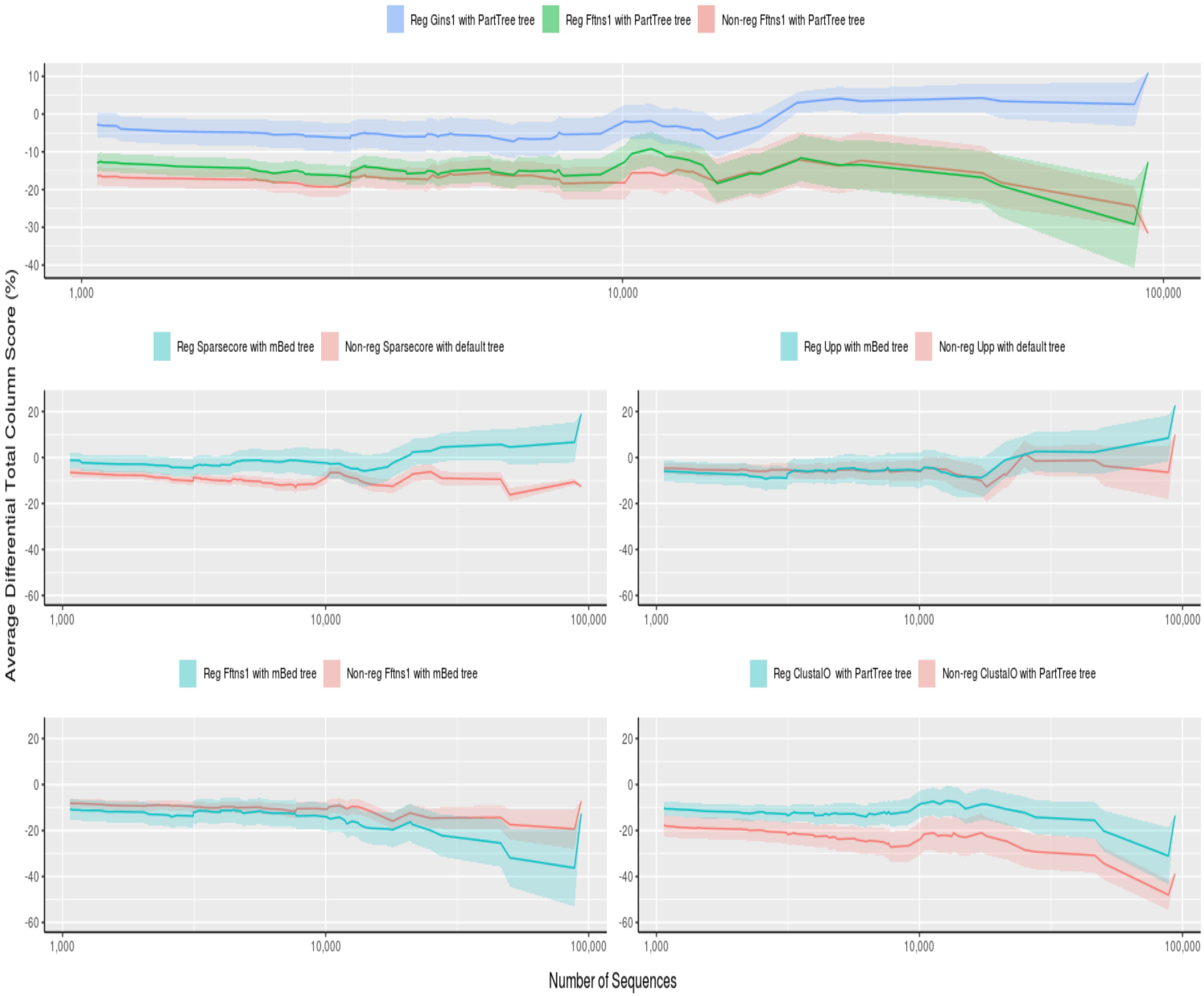

Reg Gins1 with PartTree tree
Reg Fftns1 with PartTree tree
Non-reg Fftns1 with PartTree tree
Reg Sparsecore with mBed tree
Non-reg Sparsecore with default tree
Reg Upp with mBed tree
Non-reg Upp with default tree
Reg Fftns1 with mBed tree
Non-reg Fftns1 with mBed tree
Reg ClustalO with PartTree tree
Non-reg ClustalO with PartTree tree
Average Differential Total Column Score (%)
Number of Sequences
1,000
10,000
100,000

**Fig. S1.**

**Average differential accuracy of datasets larger than Number of Sequences. (A)** The relative accuracy is defined as the difference between the TC score measured on the projection of embedded sequences and the TC score measured on the direct alignment of these same sequences with the considered method. The three alignment protocols all use a PartTree guide-trees combined with the following aligners Fftns1 in non-regressive mode (red), Fftns1 in regressive mode (green) and Gins1 in regressive mode (blue). The envelop is the standard deviation measured on the averaged values. (**B**) similar comparison between the regressive deployment of Sparsecore using a mBed guide tree (blue) and the default, non-regressive deployment of this same aligner (red). (**C**) similar comparison on UPP using a mBed guide-tree for the regressive deployment (blue) and UPP default mode for the non-regressive (red). (**D**) similar display for Fftns1 using a mBed guide-tree for the regressive (blue) and non-regressive deployment (red). (**E**) similar analysis using ClustalO as an aligner and PartTree guide-trees in regressive (blue) and non-regressive modes (red).

| A | | Over 10.000 sequences - 20 Datasets | | | | | | | |
|---|---|---|---|---|---|---|---|---|---|
| | | Sums of Pairs (SoP) | | | Total Column Score (TC) | | | CPU (s) | |
| Tree Method | MSA Algorithm | Non-regressive | Regressive | Reference | Non-regressive | Regressive | Reference | Non-regressive | Regressive |
| | | Score (%) | Score (%) | Score (%) | Score (%) | Score (%) | Score (%) | | |
| PartTree | Fftns1 | 56.34 | 59.74 | 77.10 | 29.64 | 35.16 | 47.84 | 334 | 118 |
| mBed | Fftns1 | 68.71 | 64.00 | 78.20 | 41.33 | 37.94 | 52.03 | 277 | 155 |
| PartTree | ClustalO | 50.13 | 66.58 | 78.14 | 26.94 | 42.21 | 50.54 | 3,016 | 377 |
| mBed | ClustalO | 64.97 | 71.11 | 80.54 | 39.03 | 41.91 | 53.71 | 570 | 338 |
| **Average** | | **60.04** | **65.36** | **78.50** | **34.24** | **39.31** | **51.03** | **1,049** | **247** |
| default/mBed | UPP | 71.00 | 70.71 | 80.27 | 43.80 | 44.18 | 49.88 | 8,353 | 7,186 |
| default/mBed | Sparsecore | 72.31 | 77.28 | 81.67 | 44.98 | 51.06 | 53.51 | 2,138 | 3,184 |
| PartTree | Gins1 | - | 71.51 | 78.20 | - | 47.54 | 49.46 | - | 12,477 |
| mBed | Gins1 | - | 77.14 | 81.66 | - | 50.20 | 53.07 | - | 10,833 |

| B | | Between 1,000 and 10,000 sequences - 55 Datasets | | | | | | | |
|---|---|---|---|---|---|---|---|---|---|
| | | Sums of Pairs (SoP) | | | Total Column Score (TC) | | | CPU (s) | |
| Tree Method | MSA Algorithm | Non-regressive | Regressive | Reference | Non-regressive | Regressive | Reference | Non-regressive | Regressive |
| | | Score (%) | Score (%) | Score (%) | Score (%) | Score (%) | Score (%) | | |
| PartTree | Fftns1 | 72.89 | 72.51 | 84.41 | 44.81 | 47.43 | 60.33 | 9 | 17 |
| mBed | Fftns1 | 80.77 | 78.29 | 85.93 | 56.38 | 53.75 | 63.73 | 7 | 28 |
| PartTree | ClustalO | 74.44 | 78.25 | 87.25 | 50.50 | 55.15 | 66.08 | 367 | 77 |
| mBed | ClustalO | 83.15 | 83.17 | 87.91 | 61.45 | 60.33 | 67.62 | 138 | 82 |
| **Average** | | **77.81** | **78.06** | **86.38** | **53.29** | **54.16** | **64.44** | **130** | **51** |
| default/mBed | UPP | 80.13 | 78.99 | 85.67 | 58.56 | 56.85 | 62.79 | 2,380 | 1,740 |
| default/mBed | Sparsecore | 85.24 | 86.84 | 87.69 | 60.10 | 65.10 | 65.86 | 633 | 1,091 |
| PartTree | Gins1 | - | 83.53 | 86.87 | - | 61.04 | 64.15 | - | 3,974 |
| mBed | Gins1 | - | 87.44 | 87.53 | - | 66.00 | 65.57 | - | 3,878 |

| C | | Over 92 sequences - 94 Datasets | | | | | | | |
|---|---|---|---|---|---|---|---|---|---|
| | | Sums of Pairs (SoP) | | | Total Column Score (TC) | | | CPU (s) | |
| Tree Method | MSA Algorithm | Non-regressive | Regressive | Reference | Non-regressive | Regressive | Reference | Non-regressive | Regressive |
| | | Score (%) | Score (%) | Score (%) | Score (%) | Score (%) | Score (%) | | |
| PartTree | Fftns1 | 70.95 | 71.71 | 83.64 | 44.11 | 47.13 | 59.43 | 76 | 35 |
| mBed | Fftns1 | 78.46 | 75.99 | 84.94 | 53.89 | 51.42 | 62.82 | 63 | 49 |
| PartTree | ClustalO | 71.28 | 77.89 | 85.77 | 47.81 | 55.24 | 63.66 | 858 | 126 |
| mBed | ClustalO | 80.41 | 81.73 | 86.67 | 58.25 | 58.21 | 65.19 | 203 | 121 |
| **Average** | | **75.28** | **76.83** | **85.26** | **51.02** | **53.00** | **62.78** | **300** | **83** |
| default/mBed | UPP | 78.44 | 77.99 | 85.04 | 55.36 | 54.60 | 61.06 | 3,211 | 2,590 |
| default/mBed | Sparsecore | 82.86 | 84.92 | 86.77 | 57.81 | 62.14 | 64.06 | 880 | 1,333 |
| PartTree | Gins1 | - | 81.66 | 85.21 | - | 58.89 | 61.43 | - | 5,021 |
| mBed | Gins1 | - | 85.14 | 86.68 | - | 62.36 | 63.68 | - | 4,579 |

**Table S1. Summary of SoP and TC values collected over datasets of various sizes**. (**A**) Scores measured on the 20 HomFam datasets containing more than 10,000 sequences. On each line, an Alignment algorithm is deployed in non-regressive and regressive mode using a Tree Method. Reference indicates the score of the seed alignment using the non-regressive method (note that seed alignments contain less than 10 sequences and are identically aligned by regressive and non-regressive protocols). The Central Processing Unit Time column (CPU) indicates the amount of time (seconds) used by the regressive and non-regressive protocols respectively averaged by the number of datasets. Data displayed for TC score is the same as used for Table 1 in the main text. In the SoP and TC sections, the best scoring regressive readout is highlighted in yellow and the best non-regressive is framed in red. The same layout is used on (**B**) where measures relate to the 55 datasets having between 1,000 and 10,000 sequences and (**C**) for all the 94 HomFam datasets containing 92 sequences or more.

A) Over 10,000 Sequences - TC

| Non Regressive / Regressive | MSA Algorithm | ClustalO | ClustalO | Fftns1 | Fftns1 | Sparsecore | UPP |
|---|---|---|---|---|---|---|---|
| **MSA Algorithm** | **Tree method** | **mBed** | **PartTree** | **mBed** | **PartTree** | **mBed** | **mBed** |
| ClustalO | mBed | 9.57E-02 | 7.75E-05 | 6.11E-01 | 3.11E-03 | 8.78E-01 | 9.01E-01 |
| ClustalO | PartTree | 1.31E-01 | 1.09E-04 | 2.37E-01 | 2.79E-03 | 8.35E-01 | 8.01E-01 |
| Fftns1 | mBed | 3.64E-01 | 1.58E-03 | 8.92E-01 | 1.33E-02 | 9.91E-01 | 9.53E-01 |
| Fftns1 | PartTree | 5.94E-01 | 9.93E-04 | 9.46E-01 | 6.15E-02 | 9.90E-01 | 9.78E-01 |
| Gins1 | mBed | 2.14E-03 | 8.39E-05 | 1.45E-03 | 9.08E-05 | 7.85E-02 | 9.20E-02 |
| Gins1 | PartTree | 6.04E-03 | 8.56E-04 | 8.59E-03 | 5.25E-05 | 3.74E-01 | 2.37E-01 |
| Sparsecore | mBed | 2.75E-03 | 8.39E-05 | 2.84E-03 | 6.59E-05 | 8.51E-02 | 6.31E-02 |
| UPP | mBed | 2.21E-02 | 6.05E-04 | 7.95E-02 | 4.19E-04 | 7.53E-01 | 5.09E-01 |

B) Over 10,000 Sequences - SoP

| Non Regressive / Regressive | MSA Algorithm | ClustalO | ClustalO | Fftns1 | Fftns1 | Sparsecore | UPP |
|---|---|---|---|---|---|---|---|
| **MSA Algorithm** | **Tree method** | **mBed** | **PartTree** | **mBed** | **PartTree** | **mBed** | **mBed** |
| ClustalO | mBed | 2.00E-02 | 5.57E-05 | 2.86E-01 | 6.87E-04 | 8.86E-01 | 9.17E-01 |
| ClustalO | PartTree | 2.22E-01 | 4.78E-05 | 8.00E-01 | 1.91E-02 | 9.44E-01 | 9.25E-01 |
| Fftns1 | mBed | 3.07E-01 | 1.69E-03 | 9.52E-01 | 1.20E-02 | 9.97E-01 | 9.87E-01 |
| Fftns1 | PartTree | 7.19E-01 | 2.15E-03 | 9.77E-01 | 1.80E-01 | 9.98E-01 | 9.87E-01 |
| Gins1 | mBed | 2.01E-03 | 3.18E-04 | 9.08E-04 | 7.15E-05 | 3.35E-02 | 2.32E-02 |
| Gins1 | PartTree | 2.73E-02 | 9.13E-04 | 1.90E-01 | 2.24E-04 | 5.52E-01 | 4.48E-01 |
| Sparsecore | mBed | 2.92E-03 | 2.09E-04 | 2.01E-03 | 7.15E-05 | 6.56E-02 | 1.97E-02 |
| UPP | mBed | 4.95E-02 | 9.13E-04 | 1.83E-01 | 1.58E-04 | 8.38E-01 | 5.95E-01 |

C) Between 1,000 and 10,000 Sequences - TC

| Non Regressive / Regressive | MSA Algorithm | ClustalO | ClustalO | Fftns1 | Fftns1 | Sparsecore | UPP |
|---|---|---|---|---|---|---|---|
| **MSA Algorithm** | **Tree method** | **mBed** | **PartTree** | **mBed** | **PartTree** | **mBed** | **mBed** |
| ClustalO | mBed | 6.23E-01 | 2.51E-08 | 1.11E-02 | 3.03E-08 | 8.93E-01 | 3.27E-01 |
| ClustalO | PartTree | 9.95E-01 | 7.56E-05 | 3.50E-01 | 1.55E-04 | 9.95E-01 | 8.51E-01 |
| Fftns1 | mBed | 9.99E-01 | 1.25E-02 | 6.95E-01 | 7.68E-06 | 1.00E+00 | 9.31E-01 |
| Fftns1 | PartTree | 1.00E+00 | 7.32E-01 | 9.97E-01 | 8.92E-02 | 1.00E+00 | 9.96E-01 |
| Gins1 | mBed | 6.03E-04 | 6.84E-08 | 1.31E-07 | 3.22E-10 | 8.04E-02 | 1.53E-04 |
| Gins1 | PartTree | 2.08E-01 | 5.25E-07 | 1.28E-03 | 8.25E-08 | 5.23E-01 | 7.36E-02 |
| Sparsecore | mBed | 2.64E-03 | 2.23E-07 | 6.37E-06 | 8.80E-10 | 9.36E-02 | 2.65E-03 |
| UPP | mBed | 6.27E-01 | 3.19E-03 | 8.75E-02 | 3.64E-05 | 9.30E-01 | 2.07E-01 |

D) Between 1,000 and 10,000 Sequences - SoP

| Non Regressive / Regressive | MSA Algorithm | ClustalO | ClustalO | Fftns1 | Fftns1 | Sparsecore | UPP |
|---|---|---|---|---|---|---|---|
| **MSA Algorithm** | **Tree method** | **mBed** | **PartTree** | **mBed** | **PartTree** | **mBed** | **mBed** |
| ClustalO | mBed | 4.35E-01 | 5.01E-09 | 6.84E-03 | 5.28E-09 | 9.94E-01 | 2.02E-01 |
| ClustalO | PartTree | 9.93E-01 | 2.90E-05 | 4.59E-01 | 8.15E-04 | 1.00E+00 | 8.13E-01 |
| Fftns1 | mBed | 9.99E-01 | 1.64E-03 | 7.96E-01 | 6.47E-06 | 1.00E+00 | 9.06E-01 |
| Fftns1 | PartTree | 1.00E+00 | 6.87E-01 | 9.96E-01 | 2.14E-01 | 1.00E+00 | 9.90E-01 |
| Gins1 | mBed | 1.59E-05 | 6.54E-09 | 4.51E-08 | 8.83E-11 | 2.58E-01 | 7.17E-05 |
| Gins1 | PartTree | 5.85E-02 | 2.34E-08 | 1.70E-03 | 2.23E-07 | 7.81E-01 | 5.38E-02 |
| Sparsecore | mBed | 1.12E-04 | 3.83E-08 | 1.27E-06 | 9.86E-11 | 3.16E-01 | 9.54E-04 |
| UPP | mBed | 3.67E-01 | 6.64E-03 | 1.04E-01 | 8.63E-04 | 9.97E-01 | 2.03E-01 |

E) Over 92 Sequences - TC

| Non Regressive / Regressive | MSA Algorithm | ClustalO | ClustalO | Fftns1 | Fftns1 | Sparsecore | UPP |
|---|---|---|---|---|---|---|---|
| **MSA Algorithm** | **Tree method** | **mBed** | **PartTree** | **mBed** | **PartTree** | **mBed** | **mBed** |
| ClustalO | mBed | 3.62E-01 | 3.82E-14 | 8.85E-03 | 1.69E-11 | 9.72E-01 | 4.93E-01 |
| ClustalO | PartTree | 9.44E-01 | 3.16E-10 | 1.01E-01 | 8.12E-08 | 9.96E-01 | 7.52E-01 |
| Fftns1 | mBed | 9.98E-01 | 1.64E-04 | 8.88E-01 | 3.98E-08 | 1.00E+00 | 9.92E-01 |
| Fftns1 | PartTree | 1.00E+00 | 6.04E-02 | 9.99E-01 | 2.68E-03 | 1.00E+00 | 9.99E-01 |
| Gins1 | mBed | 1.71E-05 | 8.34E-11 | 2.74E-11 | 6.79E-15 | 4.07E-02 | 5.76E-05 |
| Gins1 | PartTree | 1.88E-02 | 7.86E-10 | 4.28E-06 | 4.74E-12 | 5.10E-01 | 2.37E-02 |
| Sparsecore | mBed | 6.32E-05 | 1.67E-10 | 1.79E-09 | 1.38E-14 | 1.73E-02 | 2.01E-04 |
| UPP | mBed | 3.88E-01 | 4.83E-05 | 2.46E-02 | 8.83E-08 | 9.89E-01 | 3.18E-01 |

F) Over 92 Sequences - SoP

| Non Regressive / Regressive | MSA Algorithm | ClustalO | ClustalO | Fftns1 | Fftns1 | Sparsecore | UPP |
|---|---|---|---|---|---|---|---|
| **MSA Algorithm** | **Tree method** | **mBed** | **PartTree** | **mBed** | **PartTree** | **mBed** | **mBed** |
| ClustalO | mBed | 8.46E-02 | 3.07E-15 | 1.58E-03 | 1.49E-13 | 9.98E-01 | 3.61E-01 |
| ClustalO | PartTree | 9.33E-01 | 2.08E-11 | 3.10E-01 | 1.15E-06 | 1.00E+00 | 8.54E-01 |
| Fftns1 | mBed | 9.93E-01 | 4.57E-06 | 9.41E-01 | 2.78E-08 | 1.00E+00 | 9.93E-01 |
| Fftns1 | PartTree | 1.00E+00 | 3.62E-02 | 9.99E-01 | 2.57E-02 | 1.00E+00 | 9.99E-01 |
| Gins1 | mBed | 9.39E-08 | 4.56E-12 | 1.42E-12 | 8.20E-16 | 7.80E-02 | 1.84E-06 |
| Gins1 | PartTree | 4.43E-03 | 2.18E-11 | 3.01E-05 | 5.13E-12 | 7.30E-01 | 2.63E-02 |
| Sparsecore | mBed | 9.48E-07 | 1.10E-11 | 1.34E-10 | 7.02E-16 | 6.81E-02 | 1.09E-05 |
| UPP | mBed | 1.92E-01 | 5.30E-05 | 4.10E-02 | 7.45E-07 | 9.99E-01 | 2.94E-01 |

| < 0.001 | < 0.01 | < 0.1 | < 1 |
|---|---|---|---|

p-value

**Table S2. Wilcoxon signed-rank test p-values for differences between the regressive and non-regressive readouts.** (**A**) The p-value estimates the probability of the regressive and non-regressive having the same accuracy distribution while using the alternative hypothesis of systematically higher accuracies in the regressive mode. The test was made on datasets larger than 10,000 and using the Total Column score as a readout. Non-significant cells are colored in dark red ($0.1<P<1$), low significance cells are in light green ($0.01<P<0.1$), green ($0.001<P<0.01$) and dark green ($P<0.001$). The cells with bold borders contain p-values of the difference between directly comparable protocols (i.e. analysis displayed in the same line in Table 1). Similar analyses were made using SoP as a readout (**B**), TC and SoP for datasets containing between 1,000 and 10,000 sequences (**C, D**) respectively, SoP and TC on all the 92 HomFam datasets (E) and (F).
